# Sex-specific mechanisms underlie long-term potentiation at hippocampus-nucleus accumbens synapses

**DOI:** 10.1101/2024.01.15.575709

**Authors:** Ashley E. Copenhaver, Tara A. LeGates

## Abstract

Sex differences have complicated our understanding of the neurobiological basis of many behaviors that are key for survival. As such, continued elucidation of the similarities and differences between sexes is necessary in order to gain insight into brain function and vulnerability. The connection between the hippocampus (Hipp) and nucleus accumbens (NAc) is a crucial site where modulation of neuronal activity mediates reward-related behavior. Our previous work demonstrated that long-term potentiation (LTP) of Hipp-NAc synapses is rewarding, and that mice can make learned associations between LTP of these synapses and the contextual environment in which LTP occurred. Here, we investigate sex differences in the mechanisms underlying Hipp-NAc LTP using whole-cell electrophysiology and pharmacology. We found that males and females display similar magnitudes of Hipp-NAc LTP which occurs postsynaptically. However, LTP in females requires L-type voltage-gated Ca^2+^ channels (VGCC) for postsynaptic Ca^2+^ influx, while males rely on NMDA receptors (NMDAR). Additionally, females require estrogen receptor α (ERα) activity for LTP while males do not. These differential mechanisms converge as LTP in both sexes depends on CAMKII activity and occurs independently of dopamine-1 receptor (D1R) activation. Our results have elucidated sex-specific molecular mechanisms for LTP in an integral excitatory pathway that mediates reward-related behaviors, emphasizing the importance of considering sex as a variable in mechanistic studies. Continued characterization of sex-specific mechanisms underlying plasticity will offer novel insight into the neurophysiological basis of behavior, with significant implications for understanding how diverse processes mediate behavior and contribute to vulnerability to developing psychiatric disorders.

**SIGNIFICANCE STATEMENT:** Strengthening of Hipp-NAc synapses drives reward-related behaviors. Male and female mice have similar magnitudes of long-term potentiation (LTP) and both sexes have a predicted postsynaptic locus of plasticity. Despite these similarities, we illustrate here that sex-specific molecular mechanisms are used to elicit LTP. Given the bidirectional relationship between Hipp-NAc synaptic strength in mediating reward-related behaviors, the use of distinct molecular mechanisms may explain sex differences observed in stress susceptibility or response to rewarding stimuli. Discovery and characterization of convergent sex differences provides mechanistic insight into the sex-specific function of Hipp-NAc circuitry and has widespread implications for circuits mediating learning and reward-related behavior.

## INTRODUCTION

Sex differences in reward-related behaviors are prevalent across a variety of species. For instance, humans and rodents show clear sex differences in sensitivity to rewarding stimuli and reward value (Alarcόn et al., 2017; Aubry et al., 2022; Becker, 2016; Cullity et al., 2021; Holly et al., 2012; Legget et al., 2018; Sinclair et al., 2017; Warthen et al., 2020; Westbrook et al., 2018; Yararbas et al., 2010). There are also well-documented sex differences in the presentation of major depressive disorder in humans (Brody et al., 2018; Huang et al., 2019; Marcus et al., 2005) and depressive-like behaviors in preclinical rodent models (Baratta et al., 2019; Burke et al., 2016; Dalla et al., 2008; Goodwill et al., 2019; Liu et al., 2019; Pitzer et al., 2022; Song et al., 2018; Trainor et al., 2011; Williams et al., 2020), and males and females tend to respond differently to antidepressant treatment (reviewed in LeGates et al., 2019). Potentially underlying these behavioral differences are clear sex differences depression-related neuronal activity in humans and preclinical models (Bangasser & Cuarenta, 2021; Wang et al., 2023). Although the specific circuits and brain regions that contribute to reward-related behaviors have been identified, the precise neuronal mechanisms underlying sex differences in these behaviors and circuit function remain unknown.

The nucleus accumbens (NAc) is a key node of the reward pathway that responds to rewarding stimuli (Richter et al., 2020), integrates information from various sources to mediate goal-directed behavior (Francis & Lobo, 2017; Gruber et al., 2009), and is altered in preclinical depression models (Drysdale et al., 2017; Wacker et al., 2009). The hippocampus (Hipp) provides crucial excitatory input to the NAc which drives NAc activity and conveys spatial and contextual information to guide reward-related behavior (Bagot et al., 2015; Belujon & Grace, 2008; Britt et al., 2012; Floresco et al., 2001; Gauthier & Tank, 2018; Gill & Grace, 2013; Ito et al., 2008; LeGates et al., 2018; Lind et al., 2023; O’Donnell et al., 1999; Okuyama et al., 2016; Oliva et al., 2016; Sjulson et al., 2018; Trouche et al., 2019; Williams et al., 2020; Zhou et al., 2019). Our previous work revealed that LTP of Hipp-NAc synapses drives reward-related behaviors while exposure to chronic stress results in reduced Hipp-NAc excitatory synaptic strength, abolished LTP, and a concomitant aberration in reward-related behaviors (LeGates et al., 2018). This is supported by data from human subjects showing that functional connectivity of Hipp-striatal pathways is correlated to fluctuations in positive affect due to experiential diversity (Heller et al., 2020). These findings demonstrate a key bidirectional relationship between the strength of Hipp-NAc synapses and reward-related behaviors, highlighting the Hipp-NAc pathway as a crucial component of reward circuitry.

Given numerous examples of sex differences in reward behaviors that may be impacted by the Hipp-NAc pathway, we were interested in characterizing the molecular mechanisms underlying Hipp-NAc LTP in male and female mice. We previously found that LTP at Hipp-NAc medium spiny neurons (MSN) synapses in male mice requires NMDARs, postsynaptic Ca^2+^ influx, and CAMKII activity, but occurs independent of D1R activation (LeGates et al., 2018). Here, we performed *in vitro* electrophysiology recordings with electrical stimulation and pharmacology to characterize the mechanisms underlying Hipp-NAc MSN plasticity in females and compared them to mechanisms used in males. We found that high frequency stimulation (HFS) of hippocampal axons induced LTP in females that was similar in magnitude to HFS-induced LTP in males. While LTP was supported by postsynaptic mechanisms in sexes, the source of Ca^2+^ differed: males use NMDARs while females use L-type VGCCs. We also found that ERα activity is required for LTP in females and not males. Downstream of these differences, we found that females, like males, require CAMKII activation and have no requirement for D1R activity. Taken together, these data reveal sex-specific mechanisms that produce similar LTP at Hipp-NAc synapses which may be a key factor contributing to sex differences in behavior and disorder.

## MATERIALS & METHODS

### Animals

Adult (8–10-week-old) male and female D1dra-tdTomato or C57BL/6J mice were bred in-house or purchased directly from Jackson Laboratories. Use of D1dra-tdTomato mice allowed us to identify dopamine-1-receptor and putative dopamine-2-receptor-expressing MSNs (D1-MSN and pD2-MSN): D1-MSNs expressed tdTomato while pD2-MSNs were unlabeled. Mice were housed with same-sex cage mates in a temperature- and humidity-controlled environment under a 12:12 light cycle (lights on at 07:00). All experiments were performed in accordance with the regulations set forth by the Institutional Animal Care and Use Committee at the University of Maryland, Baltimore County.

### Mouse Brain Slice Preparation

Acute parasagittal slices containing the fornix and nucleus accumbens were prepared for whole-cell patch-clamp electrophysiology. Animals were deeply anesthetized with isoflurane, decapitated, and brains were quickly dissected and submerged in ice-cold, bubbled (carbogen: 95% O_2_/5% CO_2_) *N*-methyl-D-glucamine (NMDG) recovery solution containing the following (in mM): 93 NMDG, 2.5 KCl, 1.2 NaH_2_PO_4_, 11 glucose, 25 NaHCO_3_, 1.2 MgCl_2_, and 2.4 CaCl_2,_ pH=7.3-7.4, osmolarity=300-310 mOsm. Using a vibratome (VT1000S, Leica Microsystems), parasagittal slices (400 µm) were cut in cold, oxygenated NMDG. Slices were transferred to 32-34°C NMDG for 7-12 minutes to recover and were then transferred to room-temperature artificial cerebrospinal fluid (aCSF) containing the following (in mM): 120 NaCl, 3 KCl, 1.0 NaH_2_PO_4_, 20 glucose, 25 NaHCO_3_, 1.5 MgCl_2_·7H_2_O, and 2.5 CaCl_2,_ pH=7.3-7.4. Slices were allowed to recover for 1-hour at room-temperature before beginning electrophysiological recordings.

### Whole-Cell Recordings

We performed whole-cell patch-clamp recordings using an Axopatch 200B amplifier (Axon Instruments, Molecular Devices) and a Digidata 1550B digitizer (Axon Instruments). Slices were placed in a submersion-type recording chamber and superfused with room-temperature aCSF (flow rate 0.5-1mL/min). Patch pipettes (4-8MΩ) were made from borosilicate glass (World Precision Instruments) using a Sutter Instruments P-97 model puller.

Cells were visualized using a 60x water immersion objective (Nikon Eclipse FN-1). D1R-MSNs were identified by the expression of tdTomato while putative D2R-MSNs were cells with a similar morphology that lacked expression of tdTomato.

All recordings were performed in voltage-clamp conditions from MSNs in the NAc shell. A bipolar stimulating electrode (FHC) was placed in the fornix to electrically stimulate hippocampal axons and record evoked excitatory postsynaptic currents (EPSC). For LTP experiments, patch pipettes were filled with a solution containing 130 mM K-gluconate, 5 mM KCl, 2 mM MgCl_6_-H_2_O, 10 mM HEPES, 4 mM Mg-ATP, 0.3 mM Na_2_-GTP, 10 mM Na_2_-phosphocreatine, and 1 mM EGTA; pH=7.3-7.4; osmolarity=285-295mOsm. EPSCs were recorded from paired pulses (100ms apart) performed every 10s. A 5-minute baseline EPSC recording was obtained, then high-frequency stimulation (HFS: four trains of 100Hz stimulation for 1s with 15s between trains while holding the cell at -40mV) was used to induce LTP, followed by a 30-minute recording of EPSCs. For experiments determining I-V relationship, the patch pipette solution was composed of 135 mM CsCl, 2 mM MgCl_6_-H_2_O, 10 mM HEPES, 4 mM Mg-ATP, 0.3 mM Na_2_-GTP, 10 mM Na_2_-phosphocreatine, 1 mM EGTA, 5 mM QX-314, and 100 μM spermine; pH=7.3-7.4; osmolarity=285-295mOsm. EPSCs were collected from holding potentials ranging from -80mV to +40mV to create an I-V curve. For all pharmacological experiments, drugs (APV (Tocris, 50uM), NASPM (Tocris, 20uM), nimodipine (Tocris, 3uM), KN-62 (Tocris, 3uM), SCH23390 (Tocris, 3uM), MPP dihydrochloride (Tocris, 3uM)) were superfused over the slice for at least 15 minutes prior to recording. Recordings were discarded if there was an access resistance change >20%.

### Quantification, statistical analysis, and reproducibility

Males and females were kept separate in our analyses. We found no statistically significant difference between D1R- and pD2R-MSNs, so they are plotted together unless otherwise indicated. The sample size (*n*) per condition represents the number of mice used unless otherwise indicated in the figure caption. Multiple recordings of the same condition within the same mouse were averaged and treated as one independent sample. For LTP experiments, the last five minutes of recording were used for statistical comparisons. Pairwise comparisons using the Mann Whitney u-test were used to assess experimental condition differences. Significant pairwise comparisons were reported. A *p*-value of <0.05 was considered statistically significant, where exact p-values can be found in the figure captions. All statistical analyses were performed using Graphpad Prism 9 software. Figures were created with BioRender.com.

## RESULTS

### HFS induces Hipp-NAc MSN LTP of similar magnitude in males and females

We performed whole-cell patch-clamp electrophysiology while electrically stimulating the fornix to record Hipp-evoked excitatory postsynaptic currents (EPSC) in MSNs of the NAc shell (Fig. 1a). Slices were taken from D1dra-tdTomato mice, allowing us to distinguish between dopamine-1- and putative dopamine-2-receptor-expressing MSNs (D1-MSN and pD2-MSN) based on expression of the tdTomato reporter. In response to high frequency stimulation (HFS), we observed LTP of similar magnitude in male and female mice, with no difference between D1- and pD2-MSNs (Fig. 1). Comparison of paired-pulse ratio (PPR) suggests that LTP involves postsynaptic mechanisms in both male and female mice (Fig. 1c,e). These results indicate that LTP occurs at Hipp-NAc MSN synapses similarly in males and females, with no difference in LTP magnitude or predicted locus of plasticity mechanisms between the sexes. Since there was no difference between D1- and pD2-MSNs, cells were pooled for the remainder of the data shown.

**Figure 1.**
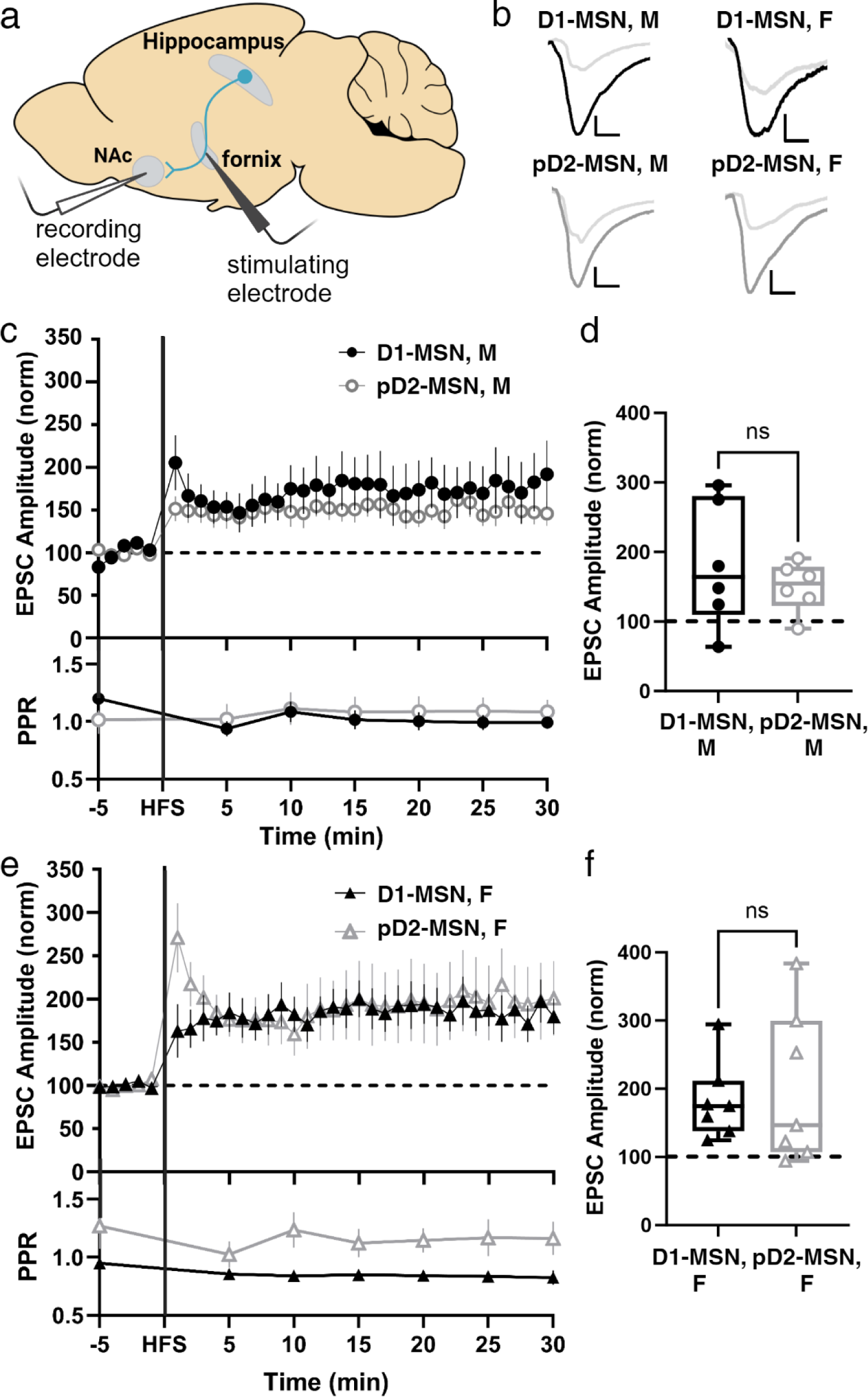
Both sexes display a similar magnitude of LTP and a predicted postsynaptic locus of plasticity. a) Recording strategy with stimulating electrode placed in the fornix and recording electrode in the NAc shell to record Hipp-evoked EPSCs from MSNs. b) Representative traces with scale bars = 40pA/10ms. c) Hippocampal-evoked EPSC amplitudes and PPR from D1- and pD2-MSNs in males. Data represent mean +/- SEM. d) Summary EPSC data from 25-30 min post-HFS revealing similar magnitudes of LTP (M D1-MSN *n*=6; M pD2-MSN *n*=6; p=0.6991, Mann-Whitney U test). e) Hippocampal-evoked EPSC amplitudes and PPR from D1- and pD2-MSNs in females. Data represent mean +/- SEM. f) Summary EPSC data from 25-30 min post-HFS revealing similar magnitudes of LTP (F D1-MSN *n*=7; F pD2-MSN *n*=7; p=0.8048, Mann-Whitney U test).

### NMDAR activity is required for male, but not female, Hipp-NAc MSN LTP

The prediction that both sexes use postsynaptic mechanisms for LTP suggests a rise in postsynaptic Ca^2+^ levels. At Hipp-NAc synapses in male mice, the source of this Ca^2+^ is NMDARs (LeGates et al., 2018). We reproduced this result in Fig. 2a,b by adding 50uM 2-Aminophosphonovaleric acid (APV), an NMDAR antagonist, to the bath. When we repeated this experiment in slices taken from female mice, we found that APV was unable to block LTP (Fig. 2c,d), suggesting the use of an NMDAR-independent pathway at female Hipp-NAc synapses. We measured AMPA:NMDA ratio at Hipp-NAc synapses in male and female mice and found no sex difference in this ratio (Fig. 2e), suggesting that there is no difference in the relative strength of the synapse. The ability to collect NMDAR-mediated currents also shows that females have NMDARs present at Hipp-NAc synapses, suggesting that the lack of requirement for NMDARs in female LTP is not explained by a lack of NMDARs in the synapse. These experiments reveal the surprising sex-specific use of an NMDAR-independent pathway for LTP at Hipp-NAc synapses.

**Figure 2.**
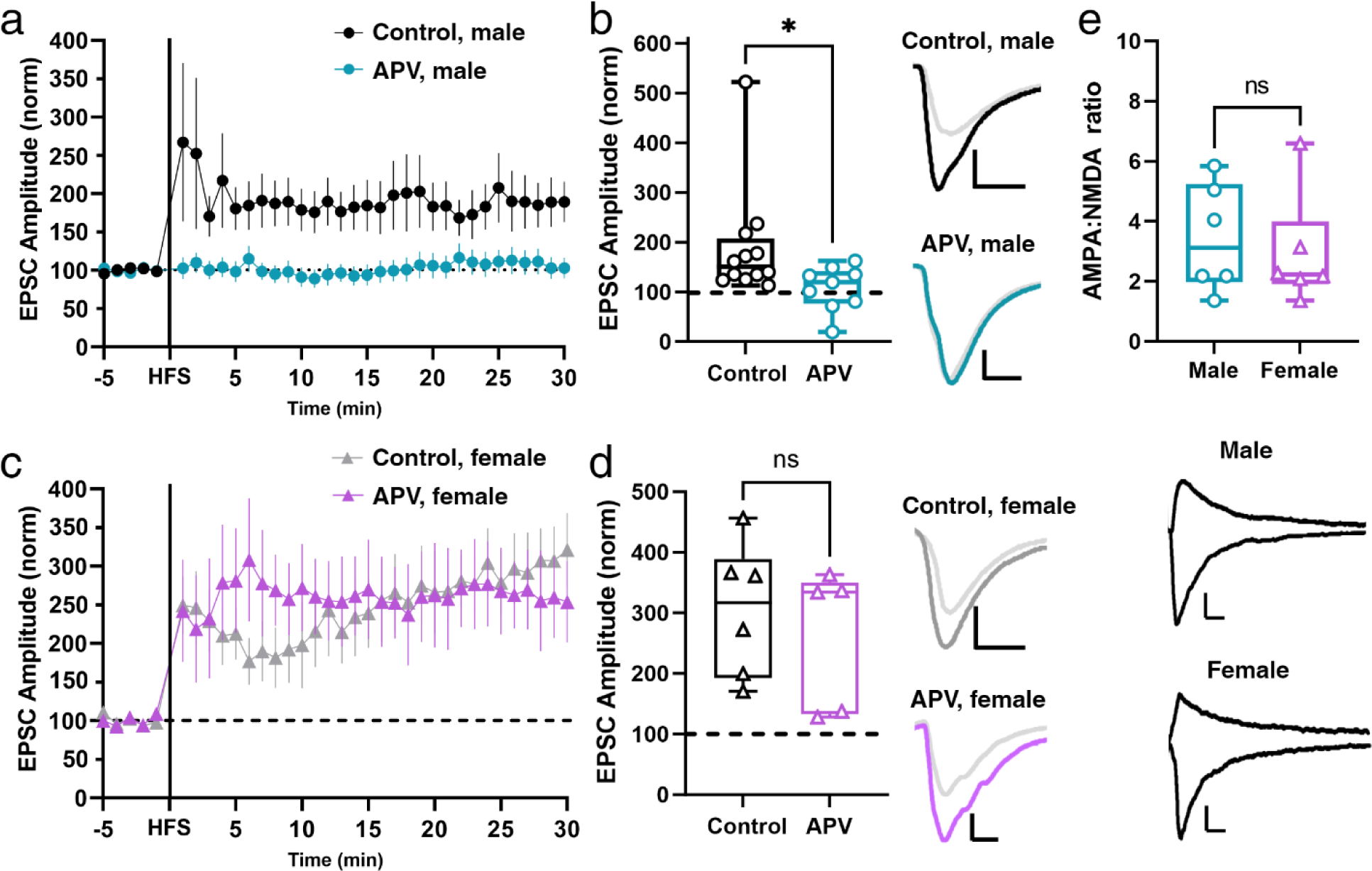
LTP is NMDAR-independent at Hipp-NAc synapses in females. a) Comparison of LTP in the presence and absence of NMDAR antagonist, APV, in males. Data represent mean +/- SEM. b) Summary EPSC data from 25-30 min post-HFS showing abolishment of LTP in APV condition (Ctrl M *n*=12; APV M *n*=9; \**p*=0.0148, Mann-Whitney U test). Representative trace scale bars = 20pA/10ms. c) Comparison of LTP in the presence and absence of NMDAR antagonist, APV, in female mice. Data represent mean +/- SEM. d) Summary EPSC data from 25-30 min post-HFS showing similar LTP magnitude in control and APV conditions (Ctrl F *n*=6; APV F *n*=5; *p*=0.4286, Mann-Whitney U test). Representative trace scale bars = 20pA/10ms. e) AMPA:NMDA ratio comparison in male and female mice reveals no sex differences in basal Hipp-NAc synaptic properties (Male *n*=6; Female *n*=6; *p*=0.7338, Mann-Whitney U test). Representative trace scale bars = 20pA/50ms.

### L-type VGCC is required for Hipp-NAc MSN LTP in females

NMDAR-independent forms of postsynaptically-expressed LTP often utilize VGCCs (Nanou & Catterall, 2018). To explore whether a postsynaptic voltage-dependent mechanism was required for LTP at Hipp-NAc synapses in females, we modified our LTP induction protocol to deliver HFS while holding the cell at -70mV rather than the depolarized (-40mV) potential we typically use. This modification effectively prevents the activation of any voltage-dependent processes during LTP induction. We found that delivering HFS in the absence of simultaneous depolarization prevented LTP induction (Fig. 3a-c), implicating the use of voltage-dependent channels for LTP in female mice.

**Figure 3.**
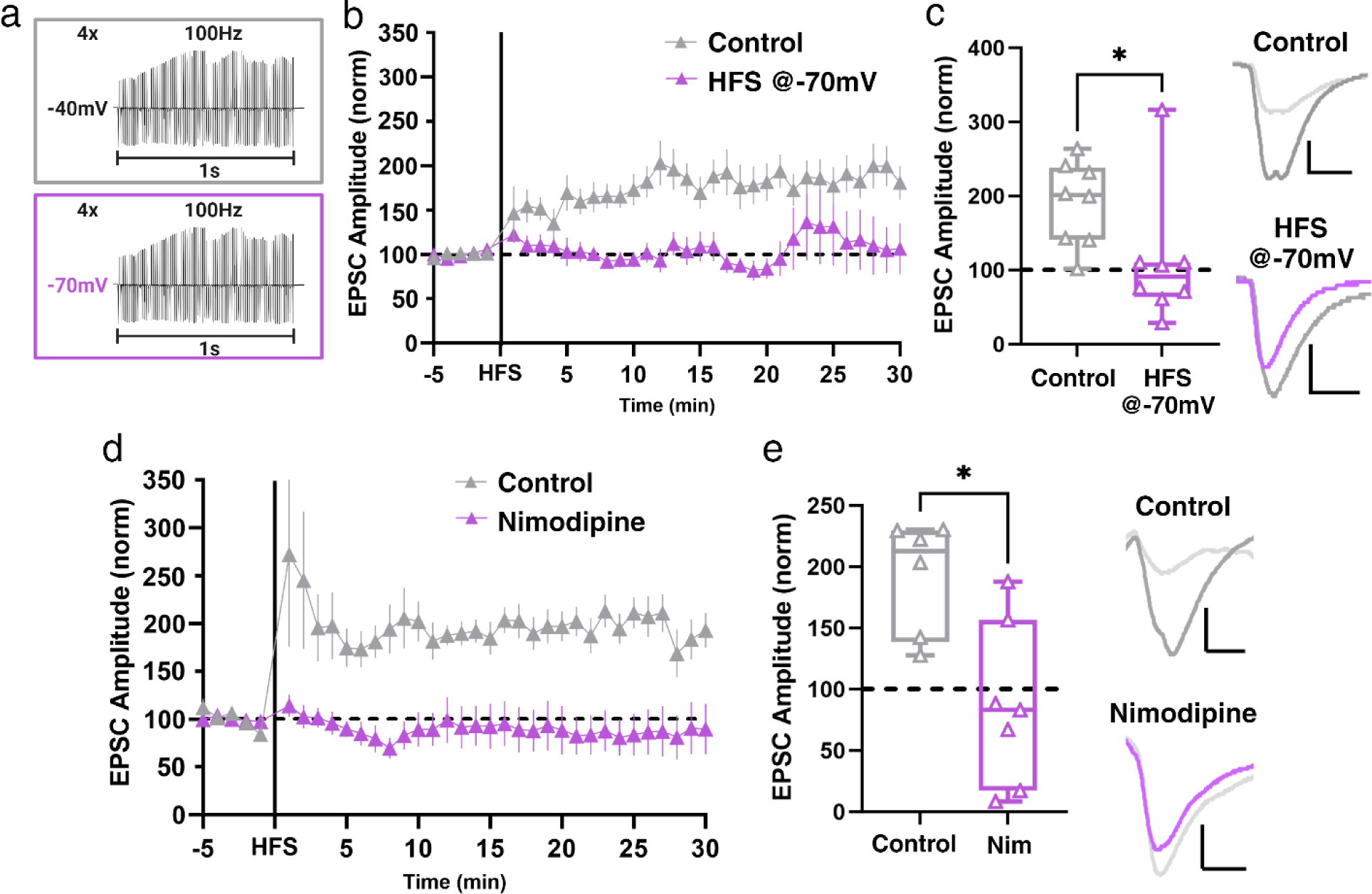
L-type VGCCs are required for LTP at Hipp-NAc synapses in females. a) Control (depolarizing cell to -40mV) and experimental (-70mV) HFS protocols. b) Comparison of LTP with control HFS and HFS @-70mV protocols. Data represent mean +/- SEM. c) Summary EPSC data from 25-30 min post-HFS showing that HFS while holding the cell at -70mV prevents LTP (Control *n*=8; HFS @-70mV *n*=8; \**p*=0.0281, Mann-Whitney U test). Representative trace scale bars = 20pA/10ms. d) Comparison of LTP in presence and absence of L-type VGCC antagonist, nimodipine (Nim). Data represent mean +/- SEM. e) Summary EPSC data from 25-30 min post-HFS reveals that Nim prevents LTP (Control *n*=6; Nim *n*=7; \**p*=0.0140, Mann-Whitney U test). Representative trace scale bars = 20pA/10ms. f) Synapse diagram showing the influx of Ca^2+^ through L-type VGCCs for LTP.

From here, we wanted to determine which type of voltage-gated channel was necessary for LTP at female Hipp-NAc synapses. L-type VGCCs have been implicated in postsynaptic forms of LTP in the amygdala and CA1 region of the hippocampus (Huber et al., 1995; Weisskopf et al., 1999) and are expressed postsynaptically within the NAc, allowing for voltage-dependent influx of Ca^2+^ into MSNs. Therefore, we hypothesized that L-type VGCCs mediate LTP at Hipp-NAc synapses in females. We tested this idea by pretreating slices with the L-type VGCC antagonist nimodipine (10uM). Bath application of nimodipine was sufficient to block LTP in female mice (Fig. 3d,e), suggesting that L-type VGCCs are required for LTP at Hipp-NAc synapses in females. Together with our recordings in the presence of APV, this demonstrates that males and females utilize distinct sources of postsynaptic Ca^2+^ to mediate LTP at Hipp-NAc synapses, where males rely on NMDARs while females require L-type VGCCs.

### CP-AMPAR are not involved in Hipp-NAc MSN LTP

To further investigate possible sex differences in postsynaptic Ca^2+^, we examined Ca^2+^-permeable AMPARs (CP-AMPAR) because they can contribute to LTP if they are present or inserted into the synapse during potentiation (Guire et al., 2008; Park et al., 2018, 2021) and they are known to contribute to multiple NAc-mediated behaviors (Carr, 2020; Jain & Woolley, 2023; Mameli et al., 2009; McCutcheon et al., 2011; Wolf & Tseng, 2012). To examine whether CP-AMPARs are present at baseline in Hipp-NAc MSN synapses, we performed a voltage-steps protocol to study the current-voltage relationship. Our data show that CP-AMPARs are not present in the Hipp-NAc MSN synapse prior to HFS (Fig. 4a). To determine whether they were preferentially inserted following HFS, we washed in 1-naphthylacetyl spermine (NASPM; 20uM), a CP-AMPAR antagonist, 10-minutes after HFS. Notably, the NASPM wash-in had no effect on EPSC amplitude (Fig. 4b,c), suggesting that insertion of CP-AMPARs does not contribute to LTP at Hipp-NAc MSN synapses in female mice. Altogether, these results rule out the involvement of CP-AMPARs in LTP at Hipp-NAc MSN synapses in females, which is similar to our previous observations in males (LeGates et al., 2018).

**Figure 4.**
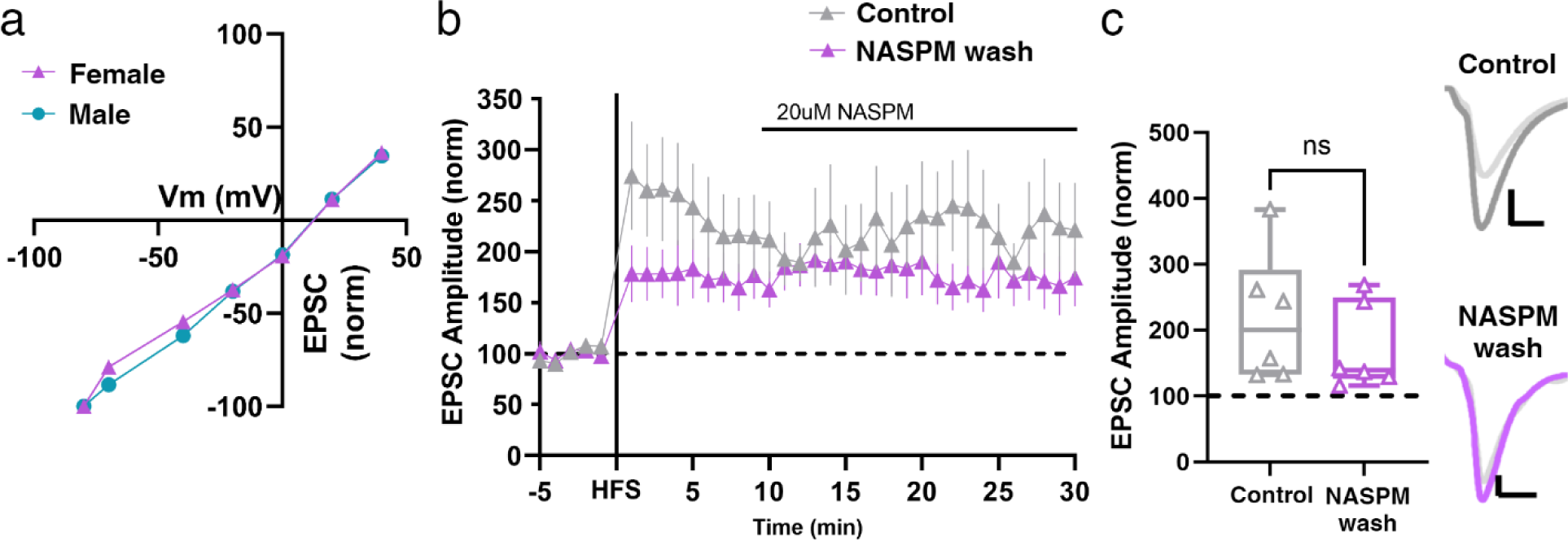
CP-AMPARs are not present at Hipp-NAc synapses and insertion of CP-AMPARs is not required for LTP in females. a) I-V relationship displays primarily AMPARs that are not permeable to Ca^2+^ at Hipp-NAc synapses (Male *n*=7 cells/4 mice; Female *n*=9 cells/3 mice). b) CP-AMPAR antagonist, NASPM, wash-on 10-minutes after HFS. Data represent mean +/- SEM. c) Summary EPSC data from 25-30 min post-HFS showing that NASPM wash has no effect on LTP (Control *n*=6; NASPM wash *n*=6; *p*=0.3939, Mann-Whitney U test). Representative trace scale bars = 40pA/10ms.

### CaMKII is required for LTP in female mice

Despite males and females differing in their sources of postsynaptic Ca^2+^, we hypothesized that the same molecular mechanisms are used downstream of this Ca^2+^ to mediate LTP. In male mice, the postsynaptic rise in Ca^2+^ initiates activation of CaMKII to cause LTP (LeGates et al., 2018). To determine whether this is consistent for LTP at Hipp-NAc synapses in females, we applied a CaMKII inhibitor (KN-62, 3uM) before recording from MSNs. We found that blocking CaMKII prevented LTP in female mice (Fig. 5), highlighting the convergence of LTP mechanisms in both sexes downstream of the Ca^2+^ channel.

**Figure 5.**
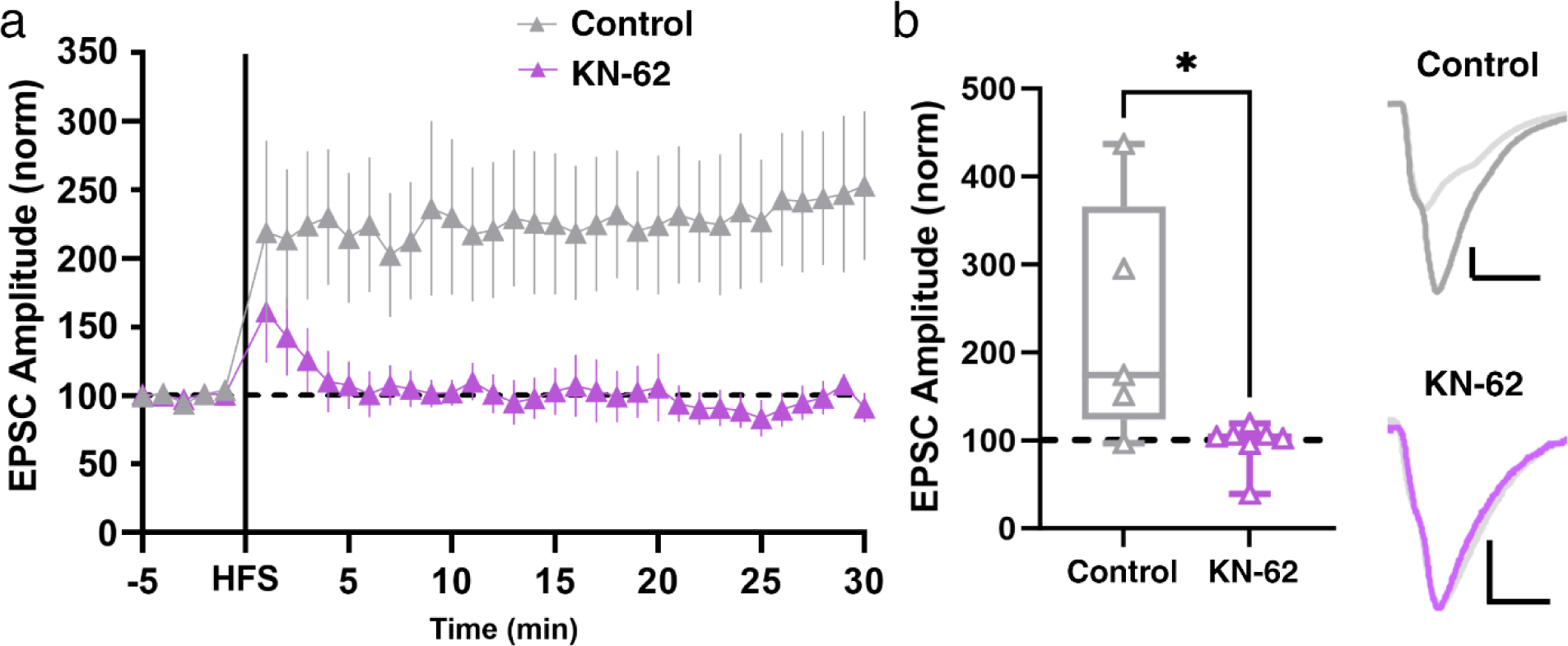
Downstream of Ca^2+^ influx, CAMKII activity is required for LTP in females. a) Comparison of LTP in the presence and absence of CAMKII antagonist, KN-62. Data represent mean +/- SEM. b) Summary EPSC data from 25-30 min post-HFS showing that KN-62 prevents LTP (Control *n*=6; KN-62 *n*=7; \**p*=0.0480, Mann-Whitney U test). Representative trace scale bars = 40pA/10ms.

### Female Hipp-NAc MSN LTP occurs independently of dopamine

Dopamine is a well-known modulator of reward-related behaviors and plays a crucial role in regulating excitatory input within the NAc (Speranza et al., 2021). Many characterized forms of LTP at excitatory synapses within the NAc require D1R activity (Du Hoffmann & Nicola, 2014; Floresco et al., 2001; Goto & Grace, 2005; Hernandez et al., 2005; Madadi Asl et al., 2019; Mameli & Lüscher, 2011; Pignatelli & Bonci, 2015; Yu et al., 2022). Interestingly, recent work has shown that Hipp-NAc LTP in male mice can still occur while dopamine receptors are blocked, demonstrating that LTP at Hipp-NAc synapses is a dopamine-independent process (LeGates et al., 2018). To test whether this was also true at Hipp-NAc synapses in females, we blocked D1R activity with the classic antagonist, SCH 23390 (3uM). Pretreatment of slices with SCH 23390 had no impact on Hipp-NAc MSN LTP in female mice (Fig. 6). This result illustrates that this dopamine-independent form of LTP in the NAc is conserved among male and female mice.

**Figure 6.**
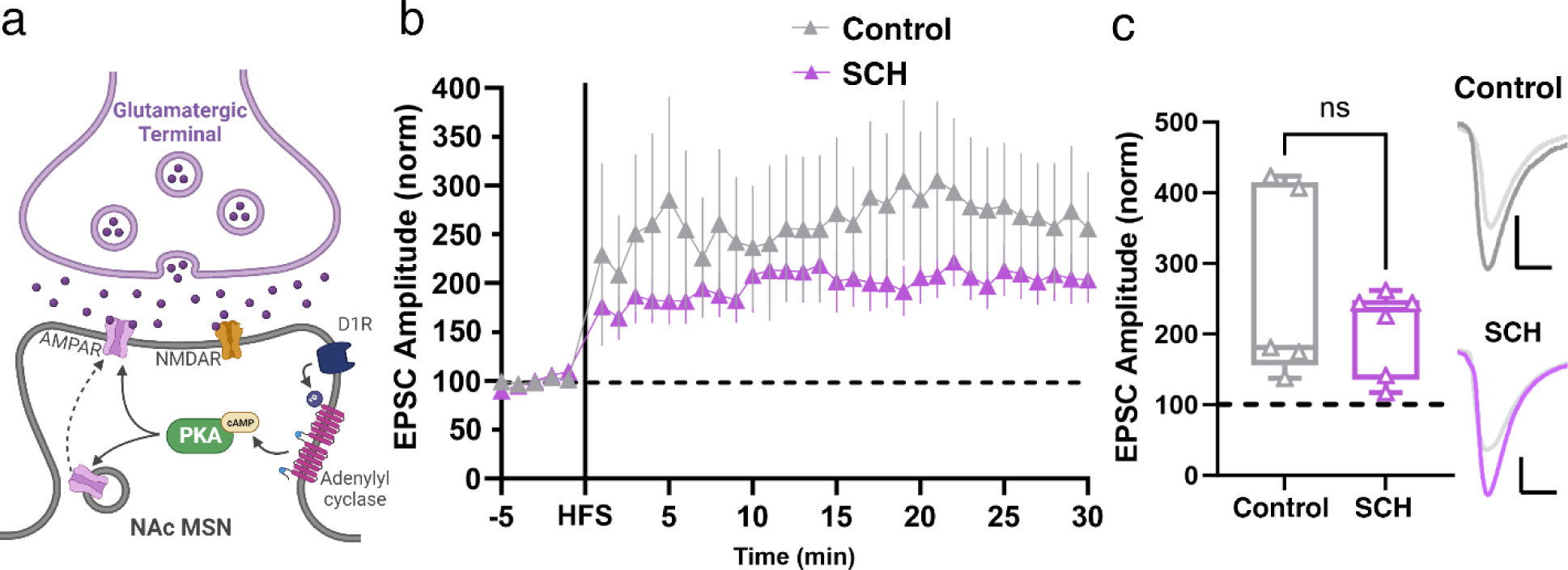
Dopamine receptor activity is not required for Hipp-NAc LTP in females. a) Schematic of D1R downstream signaling that can contribute to LTP. b) Comparison of LTP in presence and absence of D1R antagonist, SCH23390 (SCH). Data represent mean +/- SEM. c) Summary EPSC data from 25-30 min post-HFS showing that mod HFS prevents LTP (Control *n*=5; SCH *n*=6; *p*=0.7922, Mann-Whitney U test). Representative trace scale bars = 40pA/10ms.

### Estrogen receptor activity is required for female Hipp-NAc MSN LTP

Estrogen can alter excitatory synapse function and plasticity (described in detail by Frick et al., 2015; Jain & Woolley, 2023; Oberlander & Woolley, 2017) and can regulate Ca^2+^ influx via L-type VGCCs in the striatum (Mermelstein et al., 1996; Sarkar et al., 2008). Additionally, while the mechanism underlying LTP at hippocampal CA1 is similar in male and females, females have an additional requirement of membrane-bound estrogen receptor-α (ERα) activation (Gall et al., 2023; W. Wang et al., 2018). Within the NAc, ERα is expressed primarily on the membrane and has been shown to interact with GPCRs to promote other forms of plasticity (Krentzel & Meitzen, 2018; Tonn Eisinger et al., 2018). Since ERα is highly expressed in the NAc in females and has the potential to alter Ca^2+^ influx through L-type VGCCs, we postulated that ERα is required for LTP at Hipp-NAc synapses in females. We selected an ERα antagonist, MPP dihydrochloride (3uM), and added this drug to the bath composition to test the involvement of ERα in LTP at Hipp-NAc synapses. We found that pretreating slices with the ERα antagonist prevented LTP in female mice but not male mice (Fig. 7), demonstrating the sex-specific requirement of ERα activation for LTP.

**Figure 7.**
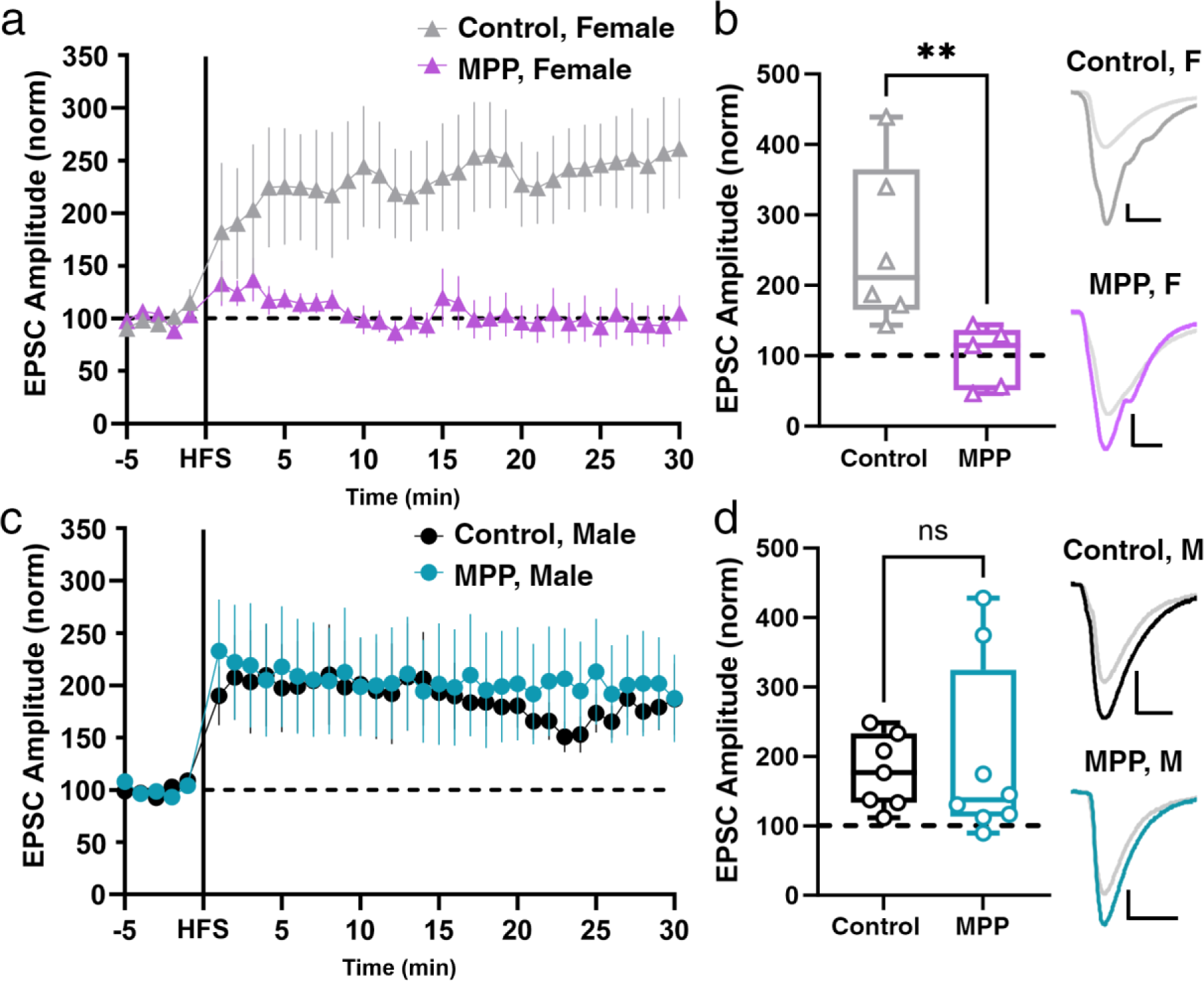
Sex-specific requirement for ERα activity for Hipp-NAc LTP. a) Comparison of LTP in the presence and absence of an ERα antagonist, MPP dihydrochloride (MPP) in female mice. Data represent mean +/- SEM. b) Summary EPSC data from 25-30 min post-HFS showing that ERα inhibition prevents LTP in female mice (Ctrl F *n*=6; MPP F *n*=5; \*\**p*=0.0087, Mann-Whitney U test). Representative trace scale bars = 20pA/10ms. c) Comparison of LTP in the presence and absence of an ERα antagonist, MPP dihydrochloride (MPP) in male mice. Data represent mean +/- SEM. d) Summary EPSC data from 25-30 min post-HFS showing that ERα inhibition has no effect on LTP in male mice (Ctrl M *n*=7; MPP M *n*=8; *p*=0.6126, Mann-Whitney U test). Representative trace scale bars = 20pA/10ms.

## DISCUSSION

Our data reveal sex-specific and -similar molecular mechanisms that underlie LTP at Hipp-MSN synapses in the NAc (Fig. 8). Both sexes display LTP of similar magnitude that relies on common mechanisms such as postsynaptic Ca^2+^ influx and CaMKII activity. However, key differences emerged when we investigated the source of the postsynaptic Ca^2+^; NMDARs are required for LTP at Hipp-NAc synapses in males, while L-type VGCCs are required in females. Furthermore, we identified a requirement for ERα in females that was not observed in males. Together, our results highlight the discovery of convergent sex differences in the molecular mechanisms underlying LTP at Hipp-NAc synapses. Given the important role for these synapses in mediating reward-related behaviors, identification of these sex differences has major implications for uncovering the neurobiological basis of sex variation in motivated behaviors and related psychiatric disorders.

**Figure 8.**
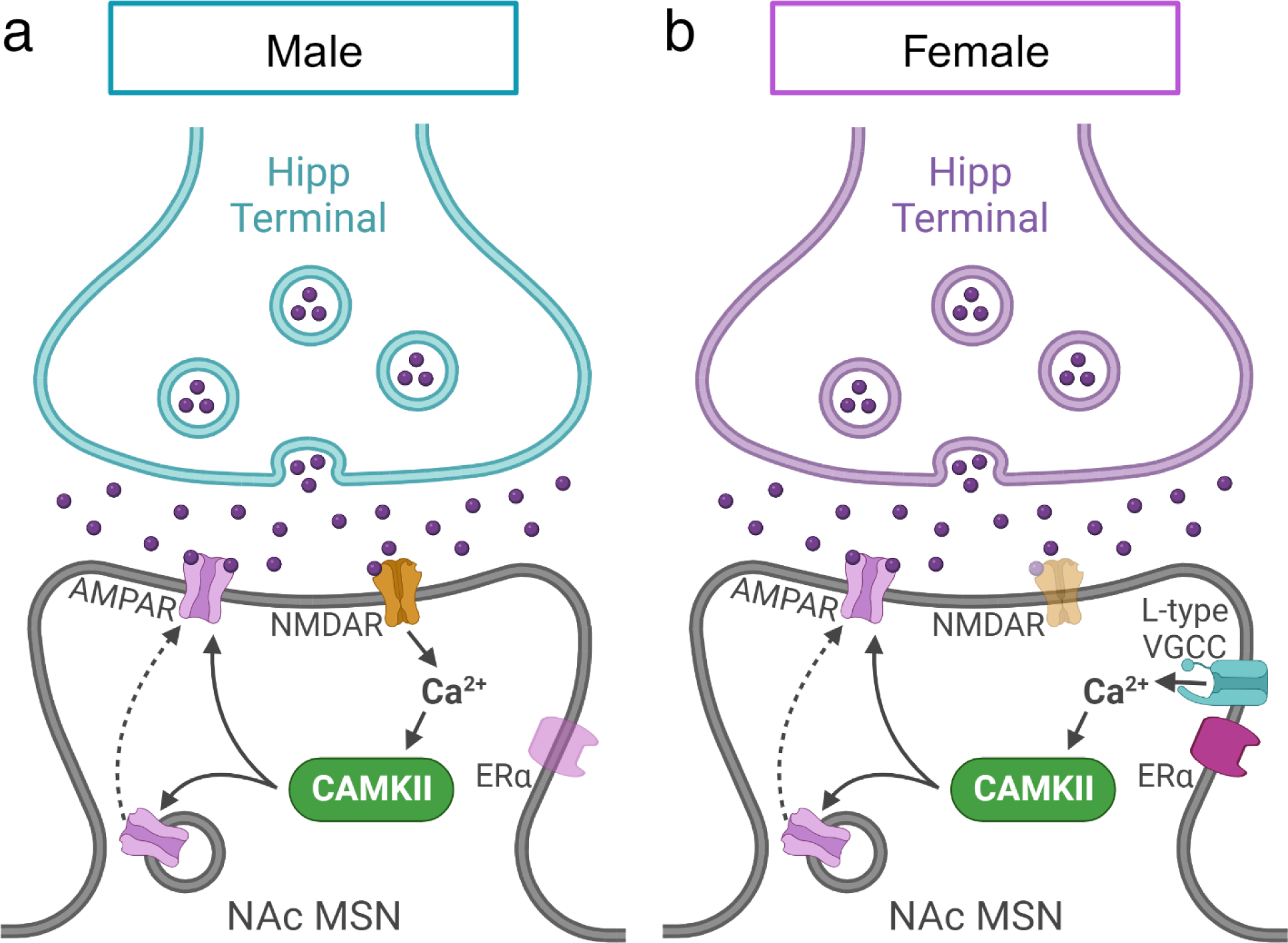
Comparison of sex-specific mechanisms involved in Hipp-NAc LTP. a) LTP at Hipp-NAc synapses in males requires NMDAR-mediated Ca^2+^ influx and CAMKII activation but does not require ERα or D1R activity. b) In females, LTP at Hipp-NAc synapses occurs with a mechanism involving L-type VGCCs instead of NMDARs for Ca^2+^ influx, CAMKII, and ERα activity, but does not require D1R activity.

### Similar LTP across sex and cell subtype

The NAc has two major subpopulations of MSNs that are classified by their dopamine receptor expression: D1-MSNs and D2-MSNs. These subpopulations differ in their neurochemical profile (Castro & Bruchas, 2019; Steiner & Gerfen, 1998) and electrophysiological properties (Al-muhtasib et al., 2018). They have been tied to different behaviors (Gagnon et al., 2017; Soares-Cunha et al., 2020), have different projection patterns (Kupchik et al., 2015; Z. Liu et al., 2022; Soares-Cunha et al., 2020), are differentially regulated by drugs of abuse (Cole et al., 2021; Gerfen et al., 1990; Smith et al., 2013), and D1-MSNs receive stronger Hipp input than D2-MSNs (Scudder et al., 2018). Despite these subtype-specific characteristics, our comparison of LTP between D1- and pD2-MSNs in males and females revealed no difference in LTP magnitude across sex or cell subtype, demonstrating similarities between sexes and subtypes in the ability to elicit LTP.

### Use of NMDAR-independent mechanism in females

NMDAR-dependent LTP is the most well-studied, prominent form of long-lasting plasticity (Bliss & Collingridge, 1993; Lüscher & Malenka, 2012), and it is the form of LTP displayed at Hipp-NAc synapses in males (LeGates et al., 2018; Fig. 2a-b). Our experiments demonstrated that NMDARs are not required for Hipp-NAc LTP in females despite sex-similar baseline electrophysiological properties, expression of AMPARs and NMDARs, and excitatory input strength onto these neurons (Willett et al., 2018; Fig. 2c-e). This is a particularly unique finding as numerous studies have shown that excitatory synapses onto NAc neurons primarily use NMDAR-dependent forms of plasticity (Floresco et al., 2001; Popescu et al., 2007; Thomas & Malenka, 2003; Vega-Villar et al., 2019), although many neurobiological studies have been performed solely in male subjects (Will et al., 2017).

NMDAR-independent mechanisms of LTP are expressed in diverse areas of the brain and can involve CP-AMPARs, metabotropic glutamate receptors, or VGCCs (Alkadhi, 2021; Bauer et al., 2002; Grover, 1998; Kullmann & Lamsa, 2008; Mameli et al., 2011; Saucier & Cain, 1995). We chose to examine L-type VGCCs because of their voltage-dependence, high expression within the NAc (Roca-Lapirot et al., 2018), and implication in LTP in other brain regions (Weisskopf et al., 1999; Huber et al., 1995). Our results showed that L-type VGCCs are required for LTP in females (Fig. 3d,e), revealing the use of sex-specific Ca^2+^ sources for LTP at Hipp-NAc synapses. Although CP-AMPARs underlie a sex-specific mechanism of synaptic potentiation in hippocampal CA1 neurons (Jain & Woolley, 2023) and are critical contributors to multiple forms of NAc plasticity that occur in response to drug exposure (Carr, 2020; Terrier et al., 2016; Wolf & Tseng, 2012), our data (LeGates et al., 2018; Fig. 4) indicated that CP-AMPARs are not involved in LTP at Hipp-NAc synapses in either sex.

L-type VGCCs and NMDARs are both voltage-dependent ion channels that allow for Ca^2+^ influx and that play critical roles in long-lasting synaptic plasticity. These channels bind calmodulin leading to Ca^2+^-dependent activation of CaMKII that is required for Hipp-NAc LTP in both sexes (LeGates et al., 2018; Ataman et al., 2007; Berger & Bartsch, 2014; Fig. 5). Despite these channel similarities, NMDARs and L-type VGCCs are differentially regulated and their dysregulation is implicated in different behaviors and diseases (Sanderson et al., 2022; Laryushkin et al., 2021; Mielnik et al., 2021; Myers et al., 2019; Ortner & Striessnig, 2015; Q. Zhou & Sheng, 2013). For example, neuronal deletion of L-type VGCCs impairs motor performance learning and increases contextual and emotional memory loss only in female mice (Klomp et al., 2022; Zanos et al., 2015). Moreover, *Cacna1c*, which encodes the Ca_v_1.2 subunit of the L-type VGCC, is a risk gene associated with multiple mood disorders (Bigos et al., 2010; Dedic et al., 2018; Jiang et al., 2023; Moon et al., 2018; Sklar et al., 2008). Increased levels of *Cacna1c* mRNA have been found in bipolar disorder postmortem brains (Bigos et al., 2010) and interactions between *Cacna1c* and sex have been identified in relation to depression (Dao et al., 2010). Additionally, *Cacna1c* haploinsufficient female mice are resilient to learned helplessness and exhibit decreased risk-taking behavior, unlike males (Dao et al., 2010), suggesting a sex-specific protective role of L-type VGCCs in regard to mood disorder-related behaviors. Given the importance of Hipp-NAc communication in mediating reward-related behaviors, the elucidation of sex-specific mechanisms underlying LTP may provide key insight to explain sex differences in behavior.

### D1R is not required for LTP at Hipp-NAc synapses

Dopamine is a critical regulator of the reward system and is typically required to induce and modulate LTP within the NAc (Floresco et al., 2001; Goto & Grace, 2005; Jay et al., 2004; Madadi Asl et al., 2019; Mameli & Lüscher, 2011; Pignatelli & Bonci, 2015; Speranza et al., 2021; Yu et al., 2022). While this occurs at many excitatory synapses in the NAc, our results (Fig. 6) and previous work (LeGates et al., 2018) show that LTP at Hipp-NAc synapses does not require D1R activation; a finding that has been observed elsewhere in the striatum and in hedonic reward learning behaviors (Berridge & Robinson, 1998; Cannon & Palmiter, 2003; Pennartz et al., 1993, p. 199). Although we have shown that D1R activation is not required for LTP at Hipp-NAc synapses, it is still possible that dopamine can alter plasticity in this pathway. In fact, other researchers have shown that dopamine can modulate the strength of LTP even when it is not required for inducing plasticity (FitzGerald et al., 2015; Otani et al., 2003; Palacios-Filardo & Mellor, 2019). Further examination of Hipp-NAc synapses in conditions with bath-applied dopamine or dopamine receptor agonists will reveal whether dopamine modulates plasticity at Hipp-NAc synapses.

### Hormones regulate Hipp-NAc synapses

Estrogen and testosterone are critical regulators of synaptic transmission and plasticity in both males and females (Barth et al., 2015; Chen et al., 2022; Lu et al., 2019; W. Wang et al., 2016; Williams et al., 2020). We show that only females require ERα activation for Hipp-NAc LTP (Fig. 7), congruent with other studies demonstrating sex differences in the requirement of estrogen receptors for LTP at hippocampal CA1 synapses (Vierk et al., 2012; W. Wang et al., 2018). The sex-specific dependence on estrogen receptor activation may stem from differences in ERα expression or function. Adult females express higher levels of membrane-localized ERα in the NAc than males (Krentzel et al., 2020; Krentzel & Meitzen, 2018). In females, membrane-localized ERα can functionally couple with metabotropic glutamate receptors (Tonn Eisinger et al., 2018) to facilitate ion flux through channels like L-type VGCCs (Subbamanda & Bhargava, 2022). Differences in ERα protein interactions could explain why we observe a sex-specific effect of ERα antagonism on LTP. Alternatively, there could be sex-specific functions associated with ERα activation. In fact, sex-specific effects of ERα activation have been reported in brain regions with no sex difference in ERα expression (Krentzel & Meitzen, 2018; Oberlander & Woolley, 2017). Further studies are necessary to identify whether ERα expression, interaction with other proteins, or downstream activated pathways are responsible for the sex-specific requirement for ERα in LTP at Hipp-NAc synapses.

### Implications of convergent sex differences in LTP at Hipp-NAc synapses

The modulation of Hipp-NAc synaptic transmission is a critical contributor to reward-related behaviors, and these synapses are altered in response to stress and cocaine (LeGates et al., 2018; Sjulson et al., 2018; Williams et al., 2020). As such, the sex differences in LTP mechanisms at Hipp-NAc synapses holds significant implications for stress- and reward-related behavior and physiology. Recent studies support the idea of sex-specific LTP mechanisms underlying sex differences in spatial learning and memory (Gall et al., 2023; Monfort et al., 2015; Safari et al., 2021; Sneider et al., 2015), so the use of sex-specific mechanisms for Hipp-NAc LTP may explain some of the sex-dependent behavioral changes that occur in response to stress and mood disorders (Brancato et al., 2017; Hodes et al., 2015; Huang et al., 2019; Salk et al., 2017; Seney & Sibille, 2014; Wei et al., 2014; Williams et al., 2020). Together, our findings highlight sex-specific mechanisms used for plasticity in a reward pathway that redefines our knowledge about LTP and offers potential molecular targets for therapeutic development to treat conditions linked to aberrant reward processing and stress.

## Acknowledgments

We would like to thank Drs. Tracy Bale, Brian Mathur, and Scott Thompson for their helpful suggestions on this project. Special thanks to Tyler Nguyen and Jennifer Pham for their assistance in maintaining the mouse colony. This work was supported by startup funds provided by UMBC and T32GM144876-02.

